# Omicron’s Intrinsic Gene-Gene Interactions Jumped Away from Earlier SARS-CoV-2 Variants and Gene Homologs between Humans and Animals

**DOI:** 10.1101/2023.02.01.526736

**Authors:** Zhengjun Zhang

## Abstract

Omicron and its subvariants have become the predominant SARS-CoV-2 variants worldwide. The Omicron’s basic reproduction number (R0) has been close to 20 or higher. However, it is not known what caused such an extremely high R0. This work aims to find an explanation for such high R0 Omicron infection. We found that Omicron’s intrinsic gene-gene interactions jumped away from earlier SARS-CoV-2 variants which can be fully described by a miniature set of genes reported in our earlier work. We found that the gene PTAFR (Platelet Activating Factor Receptor) is highly correlated with Omicron variants, and so is the gene CCNI (Cyclin I), which is conserved in chimpanzee, Rhesus monkey, dog, cow, mouse, rat, chicken, zebrafish, and frog. The combination of PTAFR and CCNI can lead to a 100% accuracy of differentiating Omicron COVID-19 infection and COVID-19 negative. We hypothesize that Omicron variants were potentially jumped from COVID-19-infected animals back to humans. In addition, there are also several other two-gene interactions that lead to 100% accuracy. Such observations can explain Omicron’s fast-spread reproduction capability as either of those two-gene interactions can lead to COVID-19 infection, i.e., multiplication of R0s leads to a much higher R0. At the genomic level, PTAFR, CCNI, and several other genes identified in this work rise to Omicron druggable targets and antiviral drugs besides the existing antiviral drugs.

## 1 Introduction

Since Omicron was first detected in Botswana in early November 2021, it has spread to become the predominant variant in circulation around the world. Compared with earlier SARS-CoV-2 variants, Omicron causes a different constellation of symptoms as well as a shorter, milder disease [1]. It can infect people who have been vaccinated or have previously had COVID-19, and even with COVID-19 vaccination protections. Scientists have tracked it in more than 120 countries but remain puzzled by a key question: where did Omicron come from [2]? Several studies have found that Omicron’s spike protein and mutations can bind or link to the ACE2 protein of turkeys, chickens, and mice/rats [3-6]. However, all things are still in the dark [1]. This paper aims to find clues to lift the darkness at genomic levels, i.e., through gene-gene interactions.

The Omicron’s basic reproduction number (R0) has been close to 20 or higher. However, it is not known what caused such an extremely high R0. This work aims to find an explanation for such high R0 Omicron infection.

Still, the pathological knowledge of the cause of all SARS-CoV-2 variants, including Omicron, and the intrinsic drivers of virus replications are unknown, at least at the genomic level and at the DNA methylation level, though many research papers have targeted these urgent needs [8-18]. Our earlier work first discovered in the literature that the genomic representation geometry spaces between SARS-CoV-2 (NP/OP PCR swabs) and COVID-19 (blood samples) are significantly different at the genomic level [16]. Using a set of optimum interactive genomic biomarkers [16], the work studying vaccine effectiveness found the adverse effects of taking BNT162b2 vaccine within the COVID-19 convalescent octogenarians [17]. Furthermore, our earlier work [18] identified COVID-19 optimum interactive DNA methylation markers.

At the genomic level, many genes have been linked to SARS-CoV-2 and COVID-19 [8-18]. The most important thing is to find reliable biomarkers. One important characteristic of reliable biomarkers is that biomarkers hold intrinsic and robust properties for different trials and cohorts. They lead to an overall accuracy being 95% or higher among all cohorts, with some cohorts being 100% accuracy. Furthermore, they are independent of extrinsic characteristics. Indeed, finding such reliable biomarkers is rather challenging. Many published gene biomarkers derived from a single trial (cohort) cannot be applied to other trials, sometimes with low efficiency. Using breast cancer diagnosis as an example, the known eight famous genes -- BRCA1, BRCA2, PALB2, BARD1, RAD51C, RAD51D, ATM were shown to be with low efficiency, see the published paper [19] and references therein. Another example is related to colorectal cancer literature. There were 56 genes identified for which gene suppression specifically inhibited the proliferation of cells harboring partial copy number loss of that gene in [43] published by ***Cell***. However, it is still not known what these genes can be truly used in cancer treatment and diagnosis as it is not clear how to use them since there are no explicit formula to follow. Our work [21] found PSMC2 and CXCL8-modulated four critical gene biomarkers for colorectal cancer can reach nearly perfect performance among seven cohort studies and high performance in a Chinese cohort study. For lung cancer, we refer readers to our earlier work [20]. These drawbacks raise outstanding concerns about many published gene biomarkers, i.e., they shouldn’t be used as biomarkers as they can mislead research in the wrong direction and mask the truth. One possible reason for the claimed biomarkers failing to be valid biomarkers may be the limitations of the analysis method and tools. A fundamental flaw is that the published gene biomarkers didn’t show their interaction with each other, and as a result, their usefulness can be rather limited.

This paper is going to apply a proven, powerful analysis approach to identify nearly perfect interactive genomic markers for COVID-19 Omicron infections [8-18].

The significance of this paper is four-fold: 1) It is the first time at the genomic level that Omicron variants’ gene-gene interactions have been discovered jumping away from earlier SARS-CoV-2 variants; 2) It is the first time that COVID-19 gene homologs between humans and animals have been discovered to be the gene CCNI; 3) It is the first time that druggable targets of Omicron infections can be different from earlier types of COVID-19 infections, and as a result, antiviral drugs for Omicron infections can have better alternative choices; 4) It is the first time that Omicron variants’ reproduction number R0 can be calculated based on gene-gene interactions which make the R0 number interpretable.

The remaining part of the paper is organized as follows. First, Section 2 briefly reviews the studying methodology. Then, Section 3 reports the data sources, analysis results, and interpretations of Omicron variants and COVID-19 infection. Next, Section 4 conducts four additional data analyses to justify the findings in Section 3. Finally, Section 5 concludes the study with discussions.

## 2 Method

We apply the newly proven method of max-linear competing logistic regression classifier to the classifications of confirmed COVID-19, healthy controls, and other COVID-19-free respiratory diseases. The new method is very different from other classical statistical and modern machine learning methods, e.g., random forest, deep learning, and support vector machine [14]. In addition, the new method has enhanced the interpretability of results, consistency, and robustness, as shown in our earlier work in studies of COVID-19 and biomarkers of several types of cancers [8-22]. This section briefly introduces the necessary notations and formulas for self-containing due to the different data structures used in this work. For continuous responses, the literature [23-24] deals with max-linear competing factor models and max-linear regressions with penalization. The max-logistic classifier has some connections to the logistic polytomous models but with different structures [25-28]. This new innovative approach can be classified as either an AI or machine learning algorithm. However, our new approach has an explicit formula and is interpretable.

Suppose *Y*_*i*_ is the *i*th individual patient’s COVID-19 status (*Y*_*i*_ = 0 for COVID-19-free, *Y*_*i*_ = 1 for infected) and 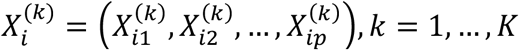 are the gene expression values, with *p* genes in this study. Here, *k* stands for the *k*th type of gene expression values drawn based on *K* different biological sampling methodologies. Note that most published works set *K* = 1, and hence the superscript (*k*) can be dropped from the predictors. In this research paper, *K* = 5, as we have five datasets analyzed in Sections 3 and 4. Using a logit link (or any monotone link functions), we can model the risk probability 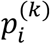 of the *i*th person’s infection status as:

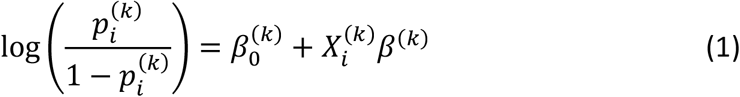

or alternatively, we write

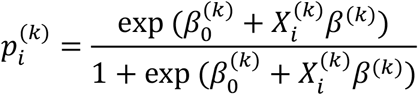

where 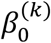 is an intercept, 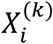 is a 1 × *p* observed vector, and *β*^(k)^ is a *p* × 1 coefficient vector which characterizes the contribution of each predictor (genes, in this study) to the risk.

Considering that there have been many variants of SARS-CoV-2 and multiple symptoms (subtypes) of COVID-19 diseases, it is natural to assume that the genomic structures of all subtypes can be different. Suppose that all subtypes of SARS-CoV-2 may be related to *G* groups of genes:

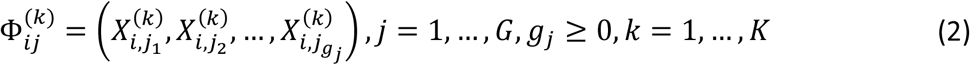

where *i* is the *i*th individual in the sample, and *g*_*j*_ is the number of genes in the *j*th group. The competing (risk) factor classifier is defined as:

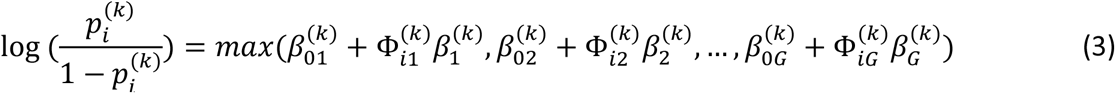

where 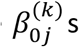 s are intercepts, 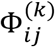 is a 1 × *g*_*j*_ observed vector, and 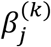 is a *g*_*j*_ × 1 coefficient vector which characterizes the contribution of each predictor in the *jth* group to the risk.

### Remark 1.

*In (3)*, 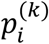 *is mainly related to the largest component* 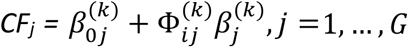, *i*.*e*., *all components compete to take the most significant effect*.

### Remark 2.

*Taking* 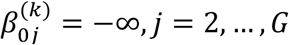, *(3) is reduced to the classical logistic regression, i*.*e*., *the classical logistic regression is a special case of the new classifier. Compared with black-box machine learning methods (e*.*g*., *random forest, deep learning (convolutional) neural networks (DNN, CNN)), and regression tree methods, each competing risk factor in (3) forms a clear, explicit, and interpretable signature with the selected genes. The number of factors corresponds to the number of signatures, i*.*e*., *G*. *This model can be a bridge between linear models and more advanced machine learning methods (black box) models. However, (3) retains the properties of interpretability, computability, predictability, and stability. Note that this remark is similar to Remark 1 in Zhang (2021) [14]*.

We have to choose a threshold probability value to decide a patient’s class label in practice. Following the general trend in literature, we set the threshold to be 0.5. As such, if 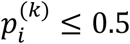, the *i*th individual is classified as being disease-free; otherwise, the individual is classified as having the disease.

With the above-established notations and the idea of a quotient correlation coefficient [29], Zhang (2021) [20] introduced a new machine learning classifier, smallest subset and smallest number of signatures (S4) as follows:

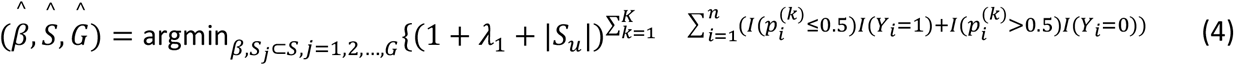

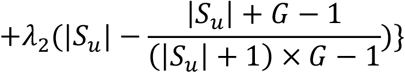

where *I*(.) is an indicative function, 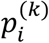 is defined in Equation (3), *S* = {1,2, …, 26369} is the index set of all genes, 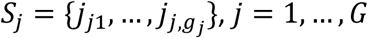 are index sets corresponding to (2), *S*_*u*_ is the union of {*S*_*j*_, *j* = 1,…, *G*}, |*S*_*u*_| is the number of elements in *S*_*u*_, *λ*_1_ ≥ 0 and *λ*_2_ ≥ 0 are penalty parameters, and 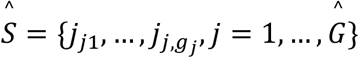 and 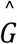 are the final gene set selected in the final classifiers and the number of final signatures.

### Remark 3.

When the S4 classifier leads to 100% accuracy, the bioequivalence and genome geometry space can be established, which is a unique property established in (4) that does not appear in other classifiers in the literature [15].

### Remark 4.

*The case of K* = 1 *corresponds to the classifier introduced by Zhang (2021) [20]. The case of K* = 1 *and λ*_2_ = 0 *corresponds to the classifier introduced by Zhang (2021) [14]*.

### Remark 5.

*All computational procedures are referred to in our earlier work [14, 20]*.

## 3 Data Descriptions, Results, and Interpretations

### 3.1 The data

The Omicron COVID-19 dataset to be analyzed in this section is publicly available at GSE201530 [30], where RNA-seq was performed with peripheral blood mononuclear cells (PBMCs) of COVID-19 patients infected by SARS-CoV-2 Omicron variant. The platform was GPL24676 Illumina NovaSeq 6000 (Homo sapiens). There are a total of 55 patients with 47 Omicron BA.2 confirmed patients and 8 healthy controls. In this study, we directly identified a small set of genes which lead to 100% accuracy among 26369 genes, and treated them as biomarkers, and compared them with our earlier findings of biomarker genes associated to earlier variants [15, 16]. In addition to this dataset, we also conduct cohort-cohort cross-validations and comparison analyses on GSE152418 [31], GSE157103 [32], GSE189039 [33], GSE205244 [34] to validate the conclusions derived from GSE201530.

### 3.2 Whether the biomarker genes for earlier SARS-CoV-2 variants are still critical

Using the reliable biomarker genes (ABCB6, KIAA1614, MND1, RIPK3, SMG1, CDC6, ZNF282, CEP72) identified for COVID-19-infected patients before Omicron variants in our earlier work directly to test whether or not these biomarker genes can still predict Omicron COVID-19 infections [14-16], we obtain the results in Table 1 (adapted from [16]).

**Table 1.** Performance of individual classifiers and combined max-competing classifiers using blood sampled data GSE201530 to classify COVID-19 infected and healthy control into their respective groups. CF-1, 2, 3, and 4 are four different classifiers. CFmax = max(CF-1,2,3,4) is the combined max-competing classifier. Raw stands for raw counts.

| Classifiers | Intercept | ABCB6 | MND1 | RIPK3 | SMG1 | CDC6 | ZNF282 | CEP72 | Accuracy | Sensitivity | Specificity |
| --- | --- | --- | --- | --- | --- | --- | --- | --- | --- | --- | --- |
| CF1(Raw) | -1.6909 |  |  |  | 0.0001 | 2.0352 | -0.6842 |  | 50.91% | 42.55% | 100% |
| CF2(Raw) | -7.5469 | -0.9264 | 5.8238 |  |  |  |  | 1.9166 | 80% | 76.60% | 100% |
| CF3(Raw) | 1.466 | 0.4688 | -1.4305 | -0.0862 |  |  |  |  | 20% | 6.38% | 100% |
| CF4(Raw) | 3.0641 | -0.8549 |  |  | 0.0001 |  | 0.6613 |  | 70.91% | 65.96% | 100% |
| CFmax |  |  |  |  |  |  |  |  | 100% | 100% | 100% |

In the table, the classifier CF1 in Equation (3) is defined as

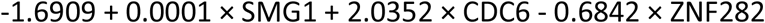

Then, 0.5 is the threshold for computing risk probability in the logistic regression function. Other classifiers are defined similarly. And CFmax is defined as the max(CF1,CF2,CF3,CF4).

We note that the number of genes in each individual classifier is determined by the algorithm which was proved to reach the smallest subset of genes, and the smallest number of classifiers. The proof appeared in [20]. In addition, the number of genes in each classifier is not necessarily three, i.e., it can be two or one, and can be four, or more. Some genes appear multiple times, while some genes just appear once.

Figure 1 presents gene expression levels and risk probabilities corresponding to different combinations in the GSE201530 dataset and Table 1.

**Figure 1.**
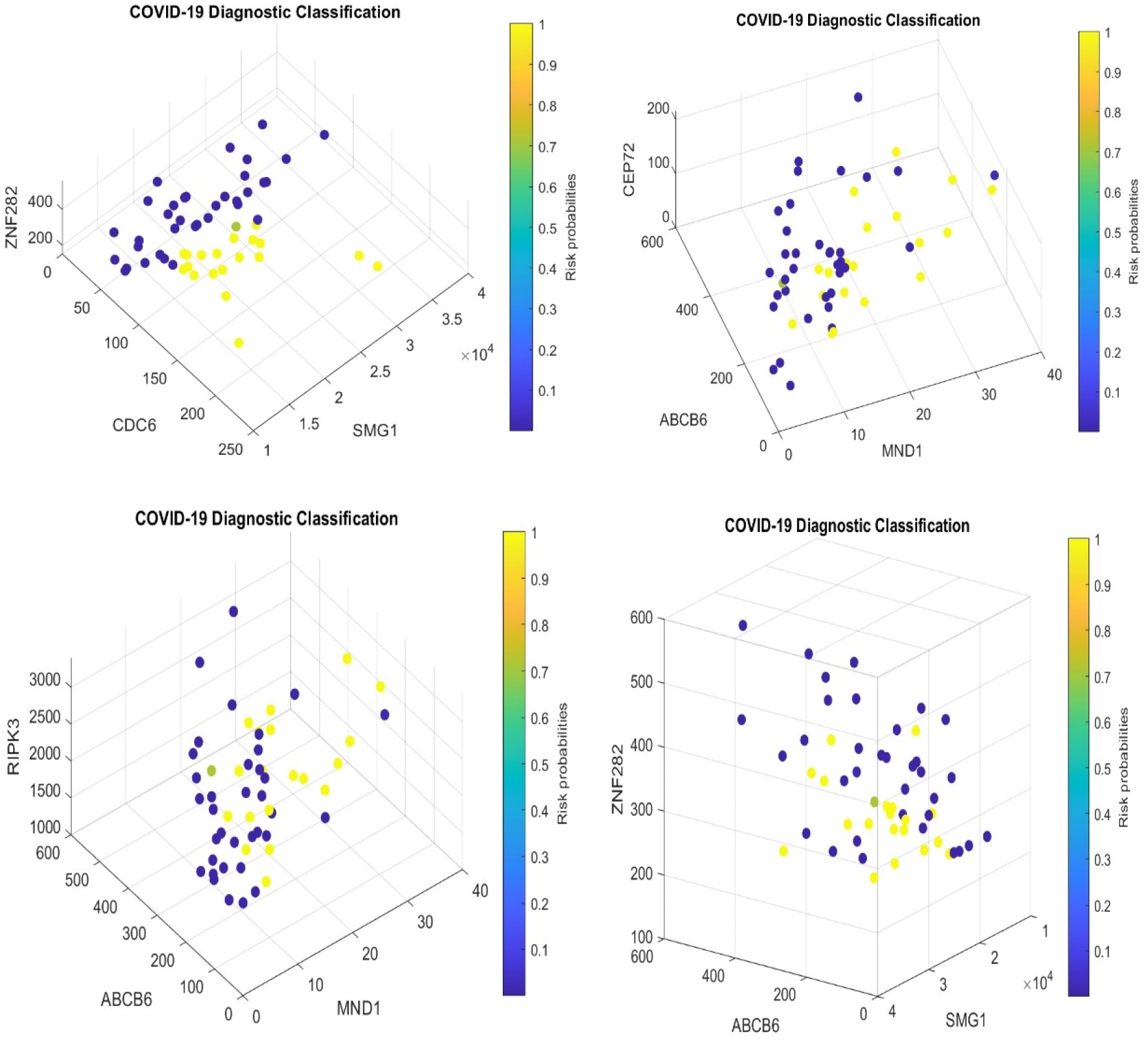
COVID-19 classifiers in Table 1: Visualization of gene-gene relationship and gene-risk probabilities. Note that 0.5 is the probability threshold. The x-/y-z-axes are gene expression levels in raw counts.

From Table 1, we see that the genes associated with the earlier variants of SARS-CoV-2 can still 100% correctly classify the patients into their respective groups using seven genes and four classifiers without considering Omicron variants’ specific features that are different from earlier variants. Compared with the earlier variants before Omicron in [16], the fitting to Omicron variants in Table 1 could be overfitted because of the constraint of genes associated with earlier variants. There are some potential issues with the fitting. First, none of the four individual classifies has an accuracy higher than 80%. Second, the patterns in Figure 1 are not clearly shown as being clustered compared to those observed in our earlier work [16] and those in Figures 2, 3 in the next section. Such observations raise questions about whether or not Omicron variants share similar gene-gene interactions with those earlier SARS-CoV-2 variants or Omicron’s intrinsic gene-gene Interactions have jumped away from earlier SARS-CoV-2 variants.

**Figure 2.**
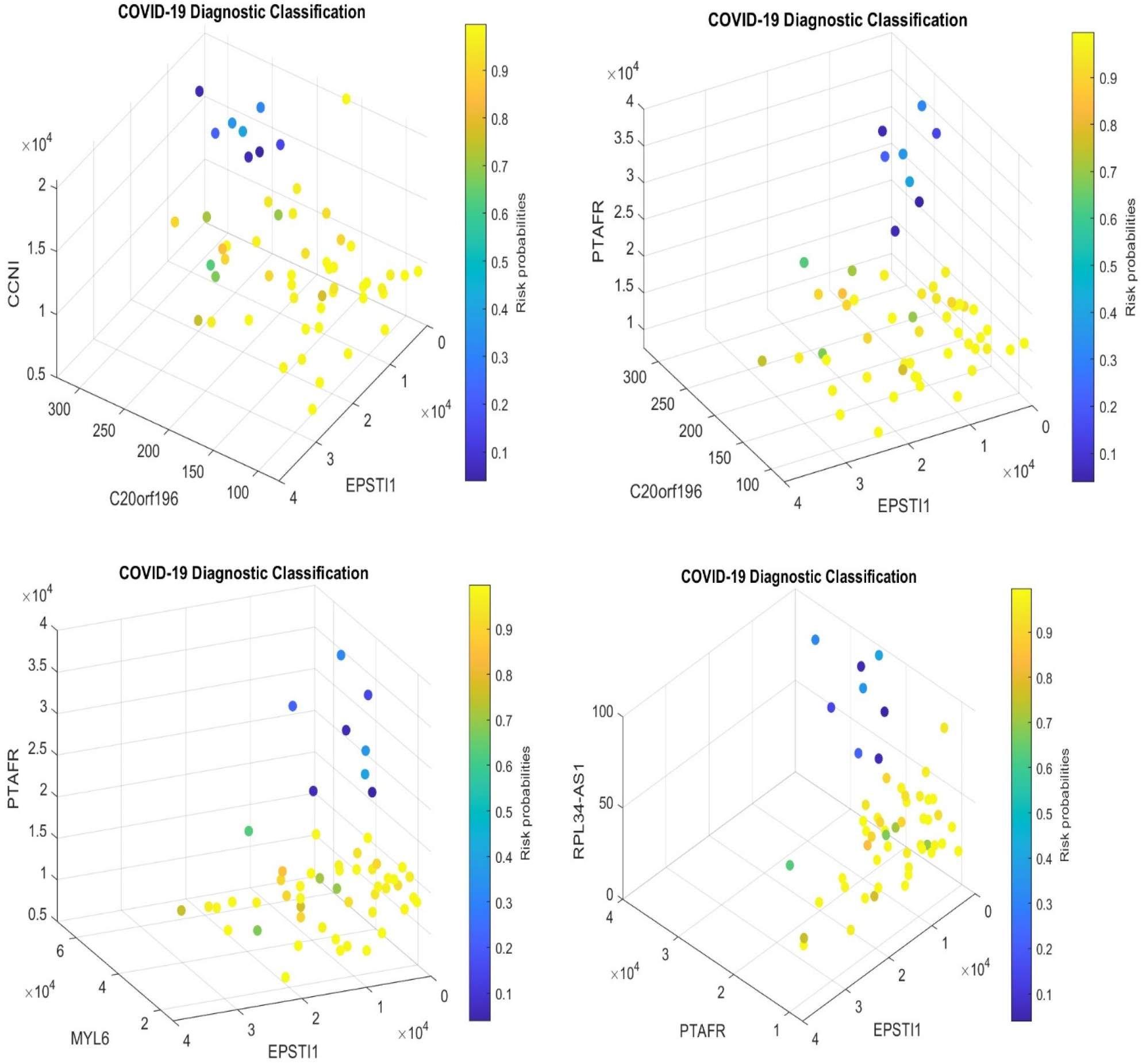
COVID-19 classifiers in Tables 3-4: Visualization of gene-gene relationship and gene-risk probabilities. Note that 0.5 is the probability threshold.

**Figure 3.**
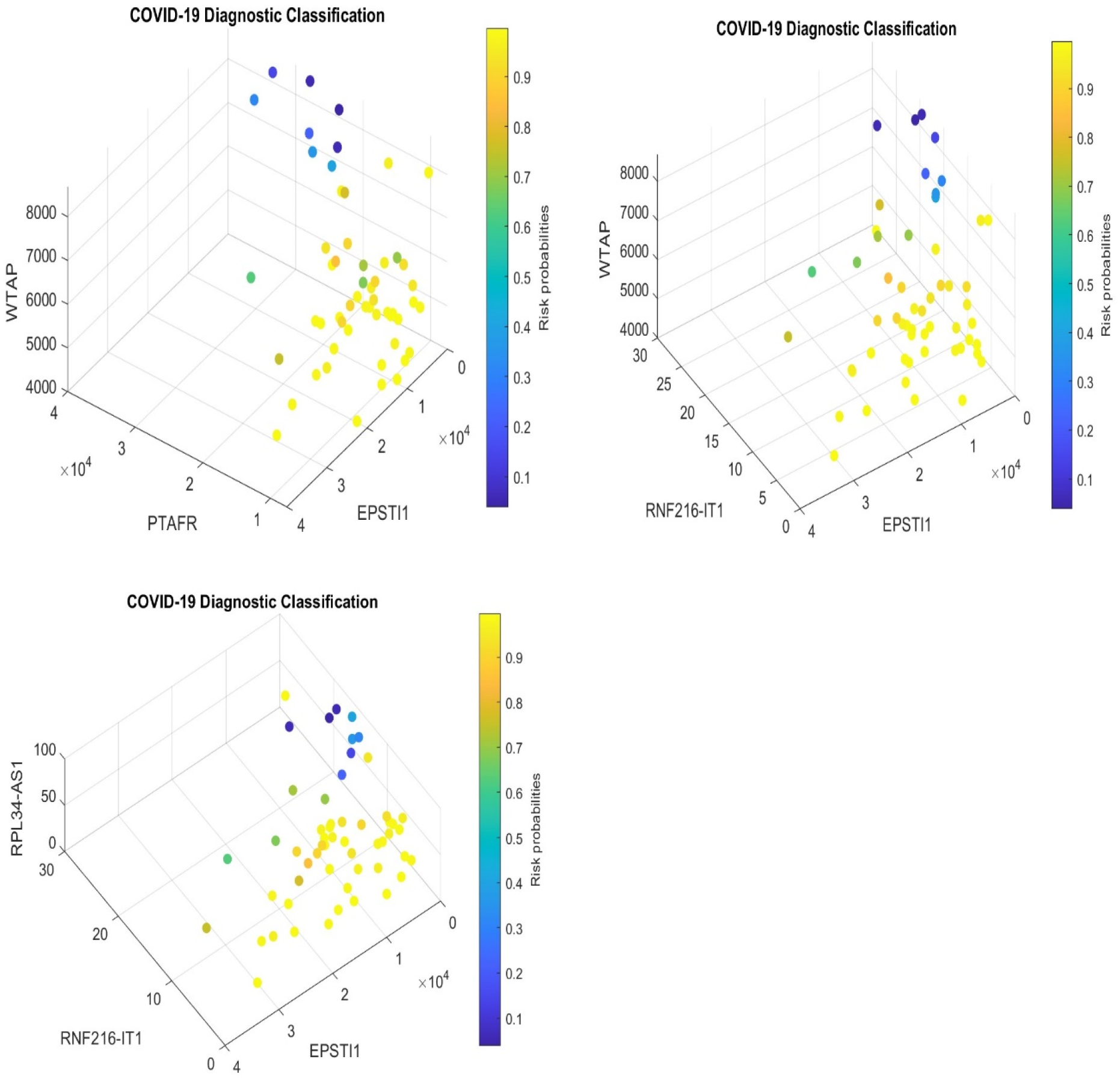
COVID-19 classifiers in Tables 3-4: Visualization of gene-gene relationship and gene-risk probabilities. Note that 0.5 is the probability threshold.

### 3.3 The biomarker genes for Omicron variants

In this section, we target Omicron variants directly using RNA-seq data from GSE201530. We found that the critical genes associated with Omicron variants are different from ABCB6, KIAA1614, MND1, RIPK3, SMG1, CDC6, ZNF282, CEP72. In addition, the gene-gene interactions in Omicron variants are simpler than those in earlier SARS-CoV-2 variants, but gene-subtype interactions are much more complex than earlier variants. Tables 2-4 report our findings.

**Table 2.** Performance of individual classifiers and combined max-competing classifiers using blood sampled data GSE201530 to classify COVID-19 infected and healthy control into their respective groups. CF-1, 2, 3, 4, 5, 6, 7 are seven different classifiers.

| gene | CF1 | CF2 | CF3 | CF4 | CF5 | CF6 | CF7 |
| --- | --- | --- | --- | --- | --- | --- | --- |
| Intercept | 10.1259 | 13.4112 | 9.4554 | 19.785 | 10.3864 | 11.3326 | 12.4117 |
| ARFGAP2 |  | -0.0015 |  |  |  |  |  |
| BTBD7 |  |  |  |  |  | -0.0025 | -0.003 |
| C20orf196 |  |  |  | -0.0439 |  |  |  |
| CCNI |  |  |  |  |  |  |  |
| DNAJB6 |  |  |  |  |  |  | -0.0011 |
| MYL6 | -0.0001 |  |  |  | -0.0001 |  |  |
| PTAFR | -0.0003 | -0.0003 | -0.0003 |  |  | -0.0003 |  |
| RNF216-IT1 |  |  |  |  |  |  |  |
| RPL34-AS1 |  |  | -0.0635 |  |  |  |  |
| ST20-AS1 |  |  |  | -0.0176 |  |  |  |
| TAGAP |  |  |  |  | -0.0003 |  |  |
| WTAP |  |  |  |  |  |  |  |
| Accuracy | 100% | 100% | 100% | 100% | 100% | 100% | 100% |
| Sensitivity | 100% | 100% | 100% | 100% | 100% | 100% | 100% |
| Specificity | 100% | 100% | 100% | 100% | 100% | 100% | 100% |

In Tables 2-4, ARFGAP2 ADP Ribosylation Factor GTPase Activating Protein 2) is a Protein Coding gene. Diseases associated with ARFGAP2 include Autoimmune Lymphoproliferative Syndrome. BTBD7 (BTB Domain Containing 7) is a Protein Coding gene. Diseases associated with BTBD7 include Skin Sarcoma. SHLD1 (Shieldin Complex Subunit 1) is a Protein Coding gene, also known as RINN3 and C20orf196, and its orthologs include mice. CCNI (Cyclin I) is a Protein Coding gene. Homologs of the CCNI gene state that the CCNI gene is conserved in humans, chimpanzee, Rhesus monkey, dog, cow, rat, chicken, zebrafish, and frog. DNAJB6 (DnaJ Heat Shock Protein Family (Hsp40) Member B6) is a Protein Coding gene. Diseases associated with DNAJB6 include Muscular Dystrophy, Limb-Girdle, Autosomal Dominant 1 and Autosomal Dominant Limb-Girdle Muscular Dystrophy. MYL6 (Myosin Light Chain 6) is a Protein Coding gene. Diseases associated with MYL6 include Noonan Syndrome 2 and Adrenal Gland Pheochromocytoma. PTAFR (Platelet Activating Factor Receptor) is a Protein Coding gene. Diseases associated with PTAFR include myringitis bullosa hemorrhagica and anxiety. RNF216-IT1 (RNF216 Intronic Transcript 1) is an RNA Gene affiliated with the lncRNA class. RPL34-AS1 is an RNA Gene, and is affiliated with the lncRNA class. ST20-AS1 (ST20 Antisense RNA 1) is an RNA Gene affiliated with the lncRNA class. Diseases associated with ST20-AS1 include exudative vitreoretinopathy 3 and exudative vitreoretinopathy. TAGAP (T Cell Activation RhoGTPase Activating Protein) is a Protein Coding gene. Diseases associated with TAGAP include Type 1 Diabetes Mellitus 21 and Febrile Seizures, Familial, 10.

WTAP (WT1 Associated Protein) is a Protein Coding gene. Diseases associated with WTAP include Wilms Tumor 1 and Wilms Tumor 5. Among its related pathways are Processing of Capped Intron-Containing Pre-mRNA and Chromatin Regulation / Acetylation. The gene information were adopted from genecards.org.

Figures 2-3 present gene expression levels and risk probabilities corresponding to different combinations in Tables 3-4.

**Table 3.**
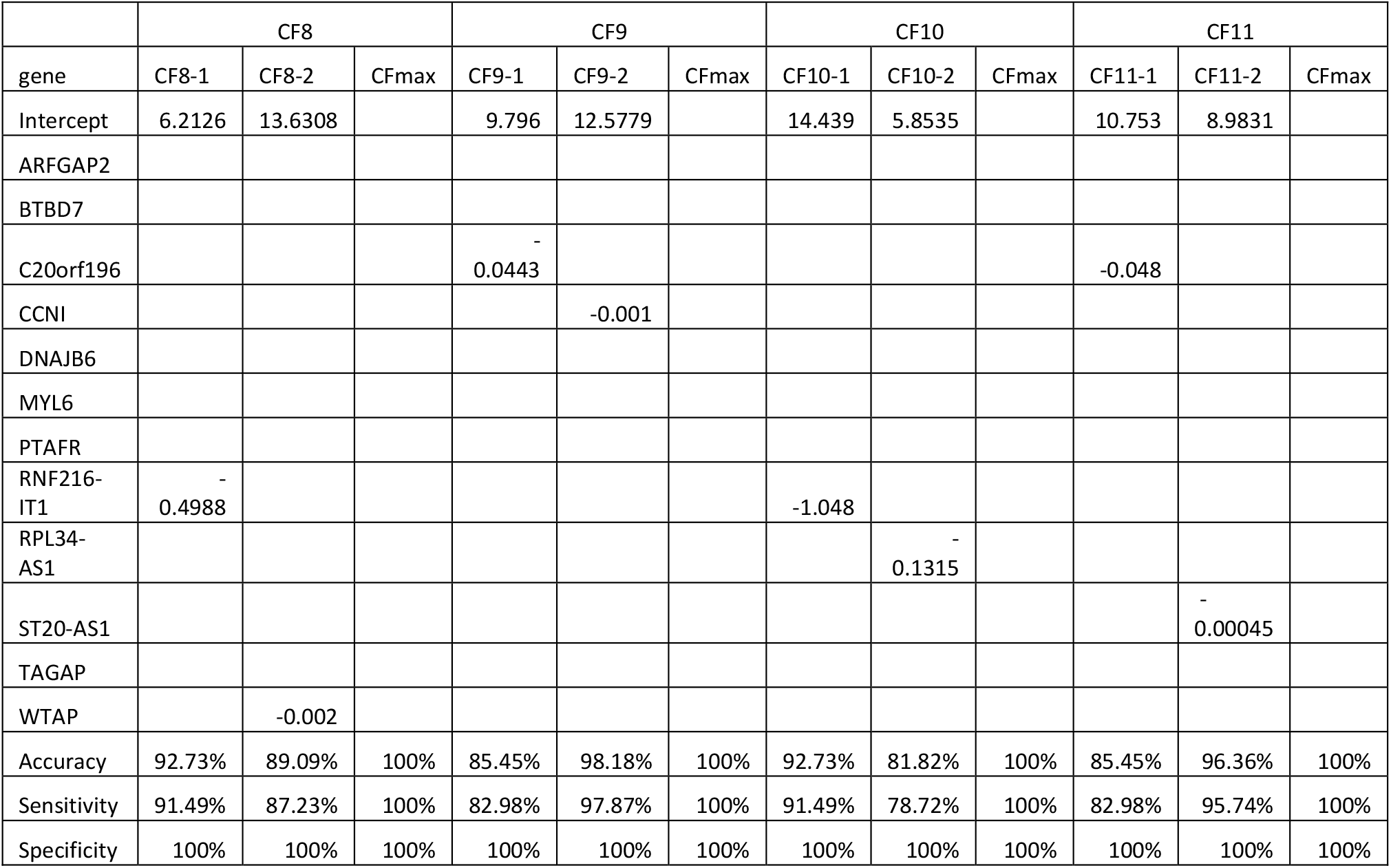
Performance of individual classifiers and combined max-competing classifiers using blood sampled data GSE201530 to classify COVID-19 infected and healthy control into their respective groups. CF8, 9, 10, 11 are four different classifiers. CFmax = max(CFi-1,2) is the combined max-competing classifier.

**Table 4.** Performance of individual classifiers and combined max-competing classifiers using blood sampled data GSE201530 to classify COVID-19 infected and healthy control into their respective groups. CF12, 13, 14 are three different classifiers. CFmax = max(CFi-1,2) is the combined max-competing classifier

|  | CF12 |  |  | CF13 |  |  | CF14 |  |  |
| --- | --- | --- | --- | --- | --- | --- | --- | --- | --- |
| gene | CF12-1 | CF12-2 | CFmax | CF13-1 | CF13-2 | CFmax | CF14-1 | CF14-2 | CFmax |
| Intercept | 8.3743 | 13.6308 |  | 8.9831 | 5.8535 |  | 7.8528 | 8.9831 |  |
| ARFGAP2 |  |  |  |  |  |  |  |  |  |
| BTBD7 |  |  |  |  |  |  |  |  |  |
| C20orf196 |  |  |  |  |  |  |  |  |  |
| CCNI |  |  |  |  |  |  |  |  |  |
| DNAJB6 |  |  |  |  |  |  |  |  |  |
| MYL6 |  |  |  |  |  |  | -0.00022 |  |  |
| PTAFR | -0.00042 |  |  | -0.00045 |  |  |  | -0.00045 |  |
| RNF216-IT1 |  |  |  |  |  |  |  |  |  |
| RPL34-AS1 |  |  |  |  | -0.13151 |  |  |  |  |
| ST20-AS1 |  |  |  |  |  |  |  |  |  |
| TAGAP |  |  |  |  |  |  |  |  |  |
| WTAP |  | -0.002 |  |  |  |  |  |  |  |
| Accuracy | 96.36% | 89.09% | 100% | 96.36% | 81.82% | 100% | 87.27% | 96.36% | 100% |
| Sensitivity | 95.74% | 87.23% | 100% | 95.74% | 78.72% | 100% | 85.11% | 95.74% | 100% |
| Specificity | 100% | 100% | 100% | 100% | 100% | 100% | 100% | 100% | 100% |

It is clear that gene-gene interactions are simpler in Omicron variants, while gene-subtype interactions in Omicron are far more complex than earlier variants [15, 16] in which interactions were analog to players interactions in a basketball team. For example, gene-gene interactions are interpreted as player-player combinations and interactions when they try to score, i.e., ball controlling strategy and ball shooting strategy. Gene-subtype interactions are interpreted as how many player combinations and ball controlling strategies a team can have. In [15,16], gene combinations involved three or more genes in each individual classifiers, while the number of combinations were three or less, which led to seven subtypes (Venn diagram) of disease classifications. Tables 2-4 and Figures 2-3 show that using two genes can classify Omicron COVID-19 infections and healthy controls in their respective groups with 100% accuracy, and we have more than 14 combinations, and more than twenty subtypes if using Venn diagram to display.

Here a subtype is defined as a unique group of patients whom can be classified by a unique set of individual classifiers among CF8 – CF14, e.g., one of CF8-1 or CF8-2, and so on. Figures 2-3 show clear patterns compared with Figure 1. The individual classifies reported in Tables 2-4 each have nearly perfect accuracy. If lower accuracies are included, or more than two genes are involved in each individual classifier, e.g., Section 3.1, more combination classifiers will lead to 100% accuracy. In the following subsections, we discuss how these observations reveal what has been missed in the literature.

In Figures 2-3, we included a gene EPSTI, which was not included in Tables 2-4. EPSTI1 (Epithelial Stromal Interaction 1) is a Protein Coding gene. Diseases associated with EPSTI1 include Lupus Erythematosus and Systemic Lupus Erythematosus (SLE). Lupus (SLE) can affect the joints, skin, kidneys, blood cells, brain, heart, and lungs, and these related symptoms have been reported in the literature studying COVID-19. EPSTI was identified as a critical gene in our earlier work [16]. In this study, we also find that the gene EPSTI has an accuracy of 72.34% in predicting Omicron COVID-19 infection. With these observations, we included the EPSTI in Figures 2-3.

We remark that the raw counts associated with the genes in Table 2 have larger ranges 10000 while those associated with the genes in Table 1 are from tens to thousands. Please refer to Figures 1-2. Such phenomenon can explain why the fitted coefficients in Table 2 are much smaller than those in Table 1. In addition, when the number of components in a combined classifier is 1, CFmax is the individual classifier itself, i.e., the individual classifier has reached the best accuracy (100% in Table 2).

To close this section, we note that if more genes are allowed, more classifiers will lead to 100% accuracy, given that a two-gene combination can lead to 100%. In our earlier work [20], the S4 classifier is defined as the miniature set of genes that lead to the best performance. In this study dataset, the number of genes in each classifier shouldn’t be more than 2. We further note that in the literature, researchers have run AI algorithms, machine learning algorithms, probability algorithms, and regular logistic regressions to find critical genes, and many genes have been reported. However, those reported genes didn’t pass cohort-to-cohort cross-validations, and as a result, their critical statements can be in doubt and potentially lead to a suboptimal direction. Nevertheless, our algorithm (4) passed cohort-to-cohort cross-validations [15-22], and as such, it deserves more attention. We note that cohort-to-cohort cross-validation is defined as that genes identified in one cohort with nearly perfect performance will perform about the same accuracy (sensitivity and specificity) among other study cohorts when directly fit them to data collected from those other cohorts.

### 3.4 Druggable targets

All twelve genes were down-regulated in their expression values after Omicron COVID-19 infections and are druggable targets. Among all twelve genes in Tables 2-4, PTAFR, CCNI, and RNF216-IT1 are the most significant genes, with 96.36%, 98.18%, and 92.73% accuracies as individual classifiers. They rise as the most druggable targets. For PTAFR, it has been discussed in the literature, e.g., Vitamin-D is known to attenuate PTAFR [35] and the reference therein; Drug– target analysis identified two receptor antagonists (rupatadine, etizolam) to PTAFR [36] and the reference therein. The diseases associated with RNF216-IT1 are not known or not reported in the literature. We will discuss CCNI in the next section. The diseases associated with other genes also deserve further investigation. In particular, diseases associated with TAGAP include Type 1 Diabetes Mellitus 21, which can be an urgent issue to investigate.

Recall that diseases associated with PTAFR include anxiety. Given Omicron variants have an extremely high R0, which has caused great public anxiety, and as a result, more people got COVID-19 infection, with many suffering from severe symptoms. Therefore, PTAFR is certainly a druggable target.

### 3.5 Omicron gene homologs between humans and animals

In the literature, the CCNI gene is conserved in chimpanzee, Rhesus monkey, dog, cow, mouse, rat, chicken, zebrafish, and frog; see e.g., [37-38] and references therein. The homologs of the CCNI gene, between humans and animals, point to a direction of lifting the dark window of why Omicron variants are so different from earlier SARS-CoV-2 variants. In Tables 1-4, and Figures 1-3, clearly, we saw that the gene-gene interactions in Omicron variants jumped away from those in earlier SARS-CoV-2 variants. More importantly, the gene CCNI has the highest individual prediction probability to predict whether or not a patient is an Omicron patient. Putting all together, it is natural to hypothesize that there is a possibility that Omicron variants were jumped from animals (very likely mice) to humans. This hypothesis deserves serious investigation through the CCNI gene. In addition, the gene C20orf196 has its orthologs including mice, which gives another genomic evidence that Omicron variants were linked to mice.

### 3.6 The Omicron reproduction number R0: a new calculation method

The Omicron’s basic reproduction number (R0) has been close to 20 or higher. In epidemiology, researchers measure R0 using contact-tracing data, and the most common method is to use cumulative incidence data. R0 values can also be simulated and estimated using ordinary differential equations. These methods can also be applied to Omicron variants. However, R0 values can do nothing to address why Omicron variants are so different from earlier SARS-CoV-2 variants. On the contrary, Omicron’s intrinsic gene-gene interactions jumped away from earlier SARS-CoV-2 variants can explain why Omicron’s R0s are so high and up to 18.86 or higher.

In Section 3.3 Tables 2-4 and Figures 2-3, as long as anyone of the fourteen individual classifiers indicates a patient’s infection status, the rest of the other thirteen classifiers will lead to the same classification. Mathematically speaking, these fourteen classifiers move together, i.e., tail co-movements. Practically speaking, a gas stove igniter can spark all fourteen burners at once. In Omicron transmissions, once Omicron viruses enter a human, as long as one of these fourteen gene-gene interactions is triggered to active, the individual is infected unless the immune system and the vaccine take effect to stop the viruses and their replications. With this structure, suppose any vulnerable individual can be infected by a specific Omicron variant through one of the fourteen (or more) ways corresponding to the fourteen classifiers in Tables 2-4, with each R0 being slightly larger than 1 (say 1.x). Then assuming the infections are independent, the overall R0 will be 1.x^14^. Suppose the overall R0 is 18.86, then we can derive an individual R0 being 1.2334. Substituting 1.2334 into Table 7 in Section 4 corresponding to the original SARS-CoV-2 virus, we get 1.2334^3^ = 1.8763, which falls into the initially estimated R0 range between 1.4 and 2.4 by the World Health Organization (WHO), and then the new calculation reflects Omicron variants.

## 4 Comparison analysis

Studies on SARS-CoV-2 and COVID-19 Infection have produced thousands of publications, with most of them at the protein level, many at the genomic level, and some at the DNA methylation level. At the genomic level, many published work simply reports the significance using simple *t*-tests or gene network analysis, e.g., [44]. However, they hardly show how gene-gene interact with each other and cohort-to-cohort cross-validations.

Our earlier work demonstrated the outperformance of our max-logistic competing risk model over AI, ML, and probability algorithms [14-22], e.g., the gene GCKR (Glucokinase Regulator) is critical for young COVID-19 patients as diseases associated with GCKR include Fasting Plasma Glucose Level Quantitative Trait Locus 5 and Maturity-Onset Diabetes of The Young, which is a severe issue in the young [18].

In this section, we evaluate the eight biomarker genes identified in our earlier work [15] and the genes reported in Section 3.2 using datasets: GSE152418, GSE157103, GSE189039, and GSE205244.

We first test the genes in Table 1 using GSE157103 and compare the results reported in [14,15]. Next, Table 5 is adapted from our earlier work [15]. Then, Table 6 presents the fitted coefficient values corresponding to the genes in Table 1 and the gene ZNF274, and related sensitivities and specificities of competing risk classifiers using TPM values. The gene ZNF274 (Zinc Finger Protein 274) is a Protein Coding gene. Diseases associated with ZNF274 include Nephrotic Syndrome, Type 4 and Immunodeficiency 21.

**Table 5.** Performance of individual classifiers and combined max-competing classifiers using blood sampled data GSE157103 to classify COVID-19 infected and other respiratory hospitalized patients into their respective groups. CF1, 2 3 are three different classifiers. CFmax = max(CFi-1,2,3) is the combined max-competing classifier. TPM stands for transcript per million, and EC stands for expected counts.

| classifiers | Intercept | ABCB6 | KIAA1614 | MND1 | RIPK3 | SMG1 | Accuracy | Sensitivity | Specificity |
| --- | --- | --- | --- | --- | --- | --- | --- | --- | --- |
| CF1 (TPM) | -0.3303 |  | 3.4153 | 0.2177 |  | -0.1248 | 69.84% | 62% | 100% |
| CF2 (TPM) | -0.7378 | -0.462 |  | 0.9093 |  | 0.0654 | 80.16% | 75% | 100% |
| CF3 (TPM) | 6.9282 |  |  |  | -0.3921 |  | 34.13% | 17% | 100% |
| CFmax |  |  |  |  |  |  | 100% | 100% | 100% |
| CF1 (EC) | -0.7877 |  | 0.0351 | 0.0181 |  | -0.0008 | 59.52% | 49% | 100% |
| CF2 (EC) | -4.6701 | -0.0408 |  | 0.2134 |  | 0.0014 | 73.02% | 66% | 100% |
| CF3 (EC) | 3.1584 |  |  |  | -0.0042 |  | 58.73% | 48% | 100% |
| CFmax |  |  |  |  |  |  | 100% | 100% | 100% |

**Table 6.**
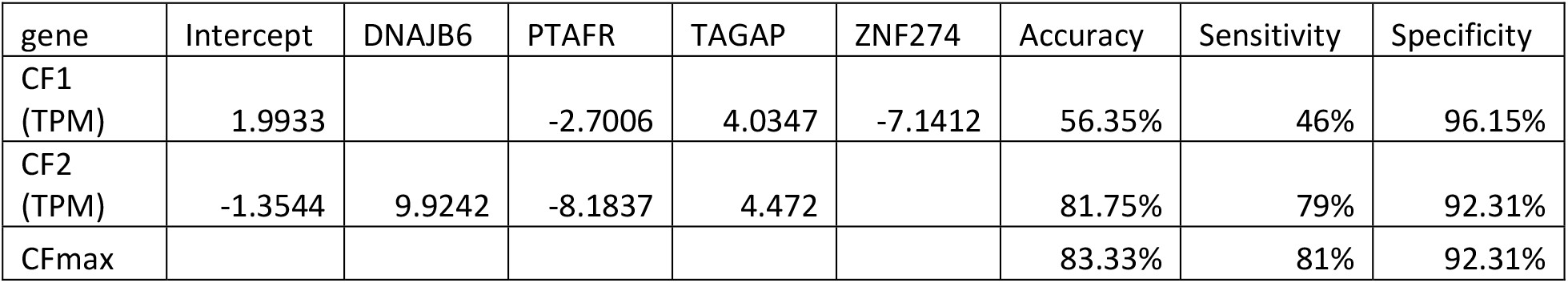
Performance of individual classifiers and combined max-competing classifiers using blood sampled data GSE157103 to classify COVID-19 infected and other respiratory hospitalized patients into their respective groups. CF1, 2 are two different classifiers. CFmax = max(CFi-1,2) is the combined max-competing classifier.

| gene | Intercept | DNAJB6 | PTAFR | TAGAP | ZNF274 | Accuracy | Sensitivity | Specificity |
| --- | --- | --- | --- | --- | --- | --- | --- | --- |
| CF1 (TPM) | 1.9933 |  | -2.7006 | 4.0347 | -7.1412 | 56.35% | 46% | 96.15% |
| CF2 (TPM) | -1.3544 | 9.9242 | -8.1837 | 4.472 |  | 81.75% | 79% | 92.31% |
| CFmax |  |  |  |  |  | 83.33% | 81% | 92.31% |

It is clear that the genes associated with Omicron variants performed badly in the original SARS-CoV-2 variants in GSE157103. We can also see that the coefficient signs of the genes DNAJB6 and TAGAP are positive in Table 6 while the corresponding coefficients in Table 2-4 are negative signs. The positive signs mean that the higher the expression values, the higher the infection risk and the higher the severity; on the contrary, the negative signs mean that the higher the expression values, the lower the infection risk and the lower the severity. These observations show that the critical gene-gene interactions and gene-subtype interactions in Omicron infections have jumped away from those in the original COVID-19 infections.

GSE152418 is an RNAseq analysis of PBMCs in a group of 17 COVID-19 subjects and 17 healthy controls. The platform is GPL24676 Illumina NovaSeq 6000 (Homo sapiens). Table 7 is adapted from our earlier work [15]. It reports the fitted coefficient values for four critical genes and related sensitivities and specificities of competing risk classifiers using raw counts. Table 8 reports the fitted coefficient values corresponding to the genes in Table 1 and the gene ZNF274, and related sensitivities and specificities of competing risk classifiers using raw counts.

**Table 7.** Performance of individual classifiers and combined max-competing classifiers using blood sampled data GSE152418 to classify COVID-19 infected and healthy control into their respective groups. CF1, 2 are two different classifiers. CFmax = max(CFi-1,2) is the combined max-competing classifier.

| Classifiers | Intercept | ABCB6 | KIAA1614 | MND1 | RIPK3 | SMG1 | Accuracy | Sensitivity | Specificity |
| --- | --- | --- | --- | --- | --- | --- | --- | --- | --- |
| CF1 (Raw) | 9.0357 |  | -0.0611 | 0.1628 |  | -0.0089 | 97.06% | 94.12% | 100% |
| CF2 (Raw) | 9.2613 | -0.2191 |  | 0.1963 |  | -0.0081 | 97.06% | 94.12% | 100% |
| CFmax |  |  |  |  |  |  | 100% | 100% | 100% |

**Table 8.**
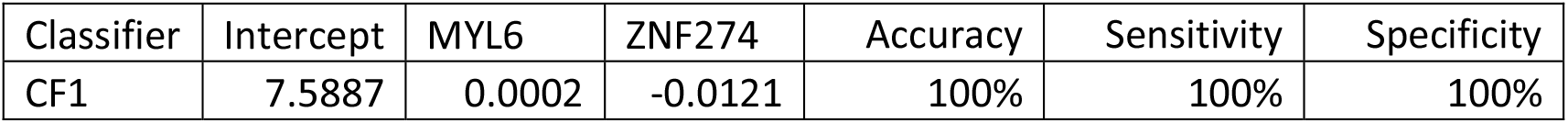
Performance of individual classifiers and combined max-competing classifiers using blood sampled data GSE152418 to classify COVID-19 infected and healthy control into their respective groups.

| Classifier | Intercept | MYL6 | ZNF274 | Accuracy | Sensitivity | Specificity |
| --- | --- | --- | --- | --- | --- | --- |
| CF1 | 7.5887 | 0.0002 | -0.0121 | 100% | 100% | 100% |

We note that both sets of genes from Table 1 and Table 2 led to 100% accuracy in Tables 7-8. The set of genes (MYL6 and ZNF274) apparently outperforms the set of genes (ABCB6, KIAA1614, MND1 and SMG1) in Tables 7-8. However, the combination of Table 5 and Table 7 together and the combination of Table 6 and Table 8 together show that the set of genes (ABCB6, KIAA1614, MND1, RIPK3, SMG1, CDC6, ZNF282, CEP72) is more informative.

The gene MYL6 in Table 8 has a positive coefficient sign, while its coefficient signs in Tables 2-4 are all negative. This phenomenon shows that even if MYL6 is connected to the original COVID-19 variants, its function in Omicron has been reversed.

GSE189039 has the overall design as RNA-seq was performed with peripheral blood mononuclear cells (PBMCs) of COVID-19 patients infected by SARS-CoV-2 Beta variant (Beta) and SARS-CoV-2 naïve vaccinated individuals. The platform was GPL24676 Illumina NovaSeq 6000 (Homo sapiens). Table 9 is adapted from our earlier work [16]. Table 10 uses the genes in Table 2.

**Table 9.** GSE189039: Characteristics of the top performed three-gene classifier CF1 for data COVID-19 vs. healthy control.

| Classifier | Intercept | ABCB6 | MND1 | CEP72 | Accuracy | Sensitivity | Specificity |
| --- | --- | --- | --- | --- | --- | --- | --- |
| CF1 | 4.742 | -0.001 | 0.0402 | -0.072 | 100% | 100% | 100% |

**Table 10.** GSE189039: Characteristics of the top performed five classifiers CF1-5 for data COVID-19 vs. healthy control.

| gene | CF1 | CF2 | CF3 | CF4 | CF5 |
| --- | --- | --- | --- | --- | --- |
| Intercept | 7.0453 | 6.203 | 6.3619 | 7.3705 | 7.9309 |
| ARFGAP2 |  | -0.0026 |  | -0.0015 |  |
| BTBD7 | -0.0026 |  |  |  | -0.0054 |
| RPL34-AS1 | -0.038 |  | -0.0495 | -0.0287 |  |
| ST20-AS1 |  |  |  |  | 0.0039 |
| WTAP |  | 0.0006 |  |  |  |
| ZNF274 |  |  | -0.0021 |  |  |
| Accuracy | 100% | 100% | 100% | 100% | 100% |
| Sensitivity | 100% | 100% | 100% | 100% | 100% |
| Specificity | 100% | 100% | 100% | 100% | 100% |

For this beta variant of SARS-CoV-2, both sets of genes from Table 1 and Table 2 led to 100% accuracy in Tables 9-10. Table 10 shows that the R0 of beta could be higher than the original R0 of COVID-19 following the discussions in Section 3.6. Also, the biomarker genes discovered for all variants are still meaningful, given they led to 100% accuracy, and these six genes in Table 10 can be more specific to the beta variants. Comparing Table 2 and Table 10, we see that the gene-gene interactions in Table 2 are different from the gene-gene interactions in Table 10. In addition, the coefficient signs of ST20-AS1 and WTAP are positive, which are different from the negative signs in Tables 2-4. These phenomena show that the gene-gene interactions in Omicron variants have jumped away from those in earlier SARS-CoV-2 variants.

Note that the most significant gene CCNI in Table 3 didn’t play any role in Tables 5-10. Given that the gene CCNI is a homologs gene between humans and animals (chimpanzee, Rhesus monkey, dog, cow, mouse, rat, chicken, zebrafish, and frog), it can be hypothesized that Omicron variants were jumped away from the animals, e.g., mouse; see also [6, 37, 38].

Next, we study the gene-gene interactions extracted from Omicron-infected patients with prior infections and without. In GSE205244, RNA-seq was performed with peripheral blood mononuclear cells (PBMCs) of COVID-19 patients infected by SARS-CoV-2 Omicron subvariants (BA.1 and BA.2). GPL24676 Illumina NovaSeq 6000 (Homo sapiens) is the platform. We test the separability of the genes in Table 1 and Table 2 in this dataset dealing with the first group (the early five days, 17 patients) and the second group (those after seven days and up to two weeks, 39 patients). Tables 11-12 report the performance of the two sets of genes.

**Table 11.** Performance of individual classifiers and combined max-competing classifiers using blood sampled data GSE205244 to classify COVID-19 Omicron infected within the early five days and after the five days into their respective groups.

| gene | CF1 | CF2 | CF3 | CFmax |
| --- | --- | --- | --- | --- |
| Intercept | 7.1559 | -2.0203 | -12.1199 |  |
| ABCB6 | -1.883 |  | -7.3861 |  |
| KIAA1614 |  | -9.1552 | 0.2633 |  |
| MND1 | -2.482 |  |  |  |
| RIPK3 |  |  | 0.8706 |  |
| CDC6 | 3.3478 |  |  |  |
| ZNF282 |  | 2.9109 |  |  |
| CEP72 |  | -9.3263 |  |  |
| Accuracy | 75% | 73.21% | 87.50% | 92.86% |
| Sensitivity | 17.65% | 11.76% | 64.71% | 82.35% |
| Specificity | 100% | 100% | 97.44% | 97.44% |

**Table 12.** Performance of individual classifiers and combined max-competing classifiers using blood sampled data GSE205244 to classify COVID-19 Omicron infected within the early five days and after the five days into their respective groups.

| Classifiers | Intercept | C20orf196 | CCNI | ST20-AS1 | TAGAP | ZNF274 | Accuracy | Sensitivity | Specificity |
| --- | --- | --- | --- | --- | --- | --- | --- | --- | --- |
| CF1 | -0.5965 | 7.4571 | -0.1026 |  |  | -0.3166 | 71.43% | 52.94% | 79.49% |
| CF2 | -0.1065 |  | -2.5866 | 9.4522 | 1.0769 |  | 78.57% | 29.41% | 100% |
| CFmax |  |  |  |  |  |  | 80.36% | 82.35% | 79.49% |

Clearly, the critical biomarker genes for earlier SARS-CoV-2 variants outperform the critical genes for Omicron variants reported in Tables 2-4. One explanation is that 19 of these 56 Omicron COVID-19 infections have previously been infected by non-Omicron variants, and the set of genes (ABCB6, KIAA1614, MND1, RIPK3, SMG1, CDC6, ZNF282, CEP72) showed 100% accuracy with Omicron infections in Table 1. Looking at Table 12, we see that the coefficient signs of CCNI are negative, which means that higher values CCNI can benefit patients to be classified into the second group (7 days after positive PCR results) which is desirable and so is ZNF274. However, the positive coefficient signs of C20orfl96, ST20-AS1, TAGAP are not the same as those in Tables 2-4, which can be explained by that all patients in GSE205244 are Omicron infected, while not all patients used in constructing Tables 2-4 (GSE205130) are.

In summary, after putting all the above analyses together, the gene CCNI is particularly functionally linked to Omicron variants. Moreover, given its homologs feature between humans and animals, together with C20orfl96, it may be safe to infer that Omicron variants jumped from animals to humans.

## 5 Discussions and Conclusions

### 5.1 Discussions

Many COVID-19 research results at the genomic level have been published in the literature. These published results explored the pathological causes of COVID-19 infection from various aspects. Due to study methodology limitations, some of the published results can hardly be cross-validated from cohort to cohort. One exception is that our earlier work [16] cross-validated thirteen genes across fourteen cohort studies with thousands of patients, heterogeneous ethics, ages, and geographical regions and showed interpretable, reliable, and robust results. Our work at the genomic level was a comprehensive study with nearly perfect performance. We didn’t find any other method that led to 100% accuracy in the literature, not even to mention interpretability. Many studies focused on only a single cohort whose representativeness cannot be assessed.

We now discuss the most significant difference between our approach and the literature approach in finding critical genes. Much attention has been paid to the individual effects of every single gene in the literature due to the study design and available analysis methods. Our approach is jointly studying gene-gene interactions and gene-subtype interactions, which were largely missed in the literature. We can see from Tables 1-12 that the effects of each gene depend on other genes in the combinations. As a result, our findings of interaction effects can be the key to the fight against COVID-19.

Many published results studied the functional effects of genes based on single gene expression value changes. They lack interaction effects study, mainly due to the limitations of study methods. As a result, they lack accuracy and may not really be useful. Using the gene CCNI as an example, in CF2 in Table 12, it must be jointly studied with another two genes ST20-AS1 and TAGAP, to fully understand its functional effects on COVID-19 as its functional effects in CF1 are different.

Since COVID-19 started in December 2019, many genes have been reported to be linked to various diseases. However, they lack mathematical proof or biological equivalence. They just happened to be significant in one cohort study. For example, SARS-CoV-2 entering the brain [39], COVID-19 vaccines complicating mammograms [40], memory loss, and ‘brain fog’ [41], and COVID-19 endothelial dysfunction can cause erectile dysfunction [42], amongst others.

Our new results show that Omicron and subvariants share gene homologs (CCNI) between human and animals, and they have mouse orthologs gene (C20orf1). We also find the earlier variants before Omicron share gene homologs (SMG1, conserved in human, chimpanzee, Rhesus monkey, dog, cow, rat, chicken, zebrafish, and frog.) The mouse orthologs gene is ABCB6. In addition, the still uncharacterized protein gene KIAA1614 is a homologs gene which is conserved in chimpanzee, Rhesus monkey, dog, cow, mouse, and rat. These significant differences again support our claims that Omicron’s Intrinsic gene-gene interactions have jumped away from earlier sars-cov-2 variants and there are gene homologs between humans and animals.

Our results are nearly perfect, with some cohort studies a 100% accuracy and others with 95% or higher accuracy. In some scenarios, such nearly perfect results can be considered too good to be true. In our earlier work [21], we argued that the traditional cross-validation method is not applicable to our model (3). Instead, we apply cohort-to-cohort cross-validation in our earlier and present work. We used a driver gene dataset to demonstrate the superiority of our model (3) compared to those algorithms built for AI, machine learning, and deep learning. We found that our results are with better precisions, and more importantly, our results are interpretable [18].

As to biochemical experiments, such tasks are beyond the scope of the paper. These work need to collaborate with a group of biochemical scientists and must be done in the most highly secured labs. Our new results light the directions of biochemical experiments, otherwise, researchers will continue explore in the dark area until there are clear evidences where Omicron came from.

### 5.2 Conclusions

In this paper, at the genomic level, we found that Omicron variants’ gene-gene interactions have been discovered jumping away from earlier SARS-CoV-2 variants. It is the first time COVID-19 gene homologs between humans and animals have been discovered to be the gene CCNI. Based on our findings, the druggable targets (CCNI, PTAFR, TAGAP, ZNF274) of Omicron infections can be different from earlier types of COVID-19 infections, and as a result, antiviral drugs for Omicron infections can have better alternative choices besides Paxlovid, Molnupiravir, and Azvudine, etc. e.g., antiviral drugs for platelet-activating factor receptor, antibodies for CCNI, drugs for Type 1 diabetes mellitus 21, and drugs for immunodeficiency. Finally, we provided a new R0 calculation method for Omicron variants, i.e., based on gene-gene interactions, which makes the R0 number interpretable.

## Acknowledgments

The authors thank insightful discussions with a group of medical doctors and scientists.

## Data Availability and Supplementary materials

The datasets are publicly available. The data links are stated in Section Data Description. Computing outputs are in a supplementary file available online https://pages.stat.wisc.edu/~zjz/OmicronJump01.zip during the review process, and the file will be submitted to the publisher after the paper has been accepted. Therefore, the results presented in this paper are all verifiable by simply checking the Excel sheets and formulas in the file.

## Competing Interests

The Authors declare no Competing Financial or Non-Financial Interests.

## Author Contributions

Zhengjun Zhang is the sole author with 100% contributions to the article.

## Statement of ethics

The authors conducted research based on published work. Therefore, the new research does not need IRB approval and a statement of ethics.

## Limitation statements

Our results are computational though we used rigorous mathematical arguments to prove biological equivalence, and the competing model with gene-gene interaction can be thought of as a revolutionary idea, and they may push a big leap in medical research. Given the nearly perfect performance, our findings demand rigorous and much deeper analysis and study in microbiology and laboratory medical tests. Although this study’s sample size is relatively small, the intrinsic relationship between SARS-CoV-2 earlier variants and Omicron variants is apparent. We demonstrated similar findings in our earlier work with additional studies [16]. The homologs of gene CCNI disclose the potential jump of Omicron variants from animals to humans demands further deep investigations. Though our results didn’t have directly biological experimental support, all findings still tell all COVID-19 problems still exist and laboratory technology are far behind to verify these findings. As such our results can still be true and meaningful and they are lights for scientific research directions and lab experiments. We believe virology experts can benefit from these findings as long as the problems cannot be solved using the classical virology knowledge.

## Notes

### Competing Interest Statement

The authors have declared no competing interest.

### Summary of Updates

The revision addressed questions and comments from readers. The author thanks the readers for their inputs.

